# A 15-Gene prognostic signature with TFAP2B functioning in Platinum Resistance of Ovarian Carcinoma

**DOI:** 10.1101/2024.09.07.611781

**Authors:** Yang Hu, Ping Wang, Jian Xiang, Lu Han, Biyun Zhang, Xiaohua Liu, Hua Nie, Guobing Chen, Weibing Qin

## Abstract

Resistance to platinum is the main challenge in the chemotherapy of ovarian cancer (OV). Therefore, developing a response signature to platinum is essential for the precision therapy of OV. Existing quantitative signatures of platinum are susceptible to batch effects and sequencing platform variations. To address this, we developed a transcriptome-based platinum signature, named PRSM, consisting of 15 genes, based on within-sample prognostic and relative expression ordering of genes, to predict individual responses to platinum in OV. The PRSM model demonstrated superior classification accuracy compared to previous quantitative signatures. Resistant samples classified by PRSM exhibited poorer overall survival, lower SNV neoantigen load, tumor mutational burden, and distinct methylation patterns compared to sensitive samples. Pathway analysis revealed the activation of MYC targets V2 and oxidative phosphorylation in resistant tumors. Single-cell analysis highlighted the roles of NK and epithelial cells in resistance. Among the 15 core genes, five (TFAP2B, KRT81, PAGE1, CRNN, UGT2B17) were linked to poor prognosis, with TFAP2B having the highest contribution to PRSM. Overexpression of TFAP2B in A2780 cells enhanced cisplatin sensitivity, while in A2780cis cells, it inhibited growth. In brief, our findings provide a multi-dimensional view of platinum resistance in ovarian cancer, introducing a robust predictive model and identifying potential therapeutic targets.

## 1. Introduction

Ovarian cancer is recognized as one of the deadliest gynecological malignancies globally, with its diagnosis often occurring at advanced stages and a high recurrence rate posing significant clinical challenges [1]. The mainstay of treatment for this formidable disease, particularly in its advanced stages, involves platinum-based chemotherapy, including agents like cisplatin and carboplatin [2, 3]. Despite the initially high response rates to these treatments, a significant portion of patients eventually develop resistance to platinum compounds, resulting in treatment failure and a dismal prognosis [3]. This resistance is thought to arise from the emergence of specialized tumor cells during treatment that are impervious to chemotherapy, leading to the progression of the disease and poor patient outcomes.

Recent research has cast light on the critical roles of genetic and epigenetic modifications, as well as variations in gene expression, in driving chemotherapy resistance in ovarian cancer [4–6]. Despite these advances, our understanding of the key genes and their regulatory networks influencing clinical outcomes is still evolving. Thus, identifying these DEGs provides critical insights into the molecular pathways affecting drug efficacy and opens avenues for developing targeted therapies and personalized treatment strategies to counteract resistance. Nevertheless, pinpointing key genes that significantly influence clinical outcomes and creating a robust risk model to predict platinum resistance are urgent needs. The complex interplay between gene expression and chemotherapy resistance remains a focal point of investigation, with the goal of deciphering the intricate genetic and epigenetic landscapes that confer resistance [4, 7, 8]. Techniques such as differential expression gene (DEG) analysis and the examination of competing endogenous RNA (ceRNA) networks and transcription factor (TF) regulatory networks have proven invaluable in identifying and comprehending these resistance mechanisms [9, 10]. Additionally, genomic variants, including Tumor Mutational Burden (TMB) [11], Single Nucleotide Variant (SNV) Neoantigens [12, 13], Homologous Recombination Deficiency (HRD) [14], and copy number variants (CNV) [15], have been linked to changes in drug metabolism and efflux, thereby affecting chemotherapy effectiveness. However, integrating these complex data to forecast clinical outcomes and guide treatment strategies remains a significant challenge, underlining the need for innovative analytical methods.

This study aims to address these gaps by conducting a comprehensive analysis of the TCGA-OV dataset to identify genes associated with platinum resistance and their impact on patient survival. We performed differential expression analysis and Cox regression to pinpoint key resistance-related genes. Based on these findings, we developed a predictive platinum resistance scoring model (PRSM) using elastic net regression. The model’s predictive accuracy was validated using receiver operating characteristic (ROC) curves and tested across multiple independent datasets to ensure robustness. Further, we explored the mutation and epigenetic landscapes associated with the identified resistance genes. Pathway analysis was conducted to identify key biological processes and signaling pathways implicated in platinum resistance. Single-cell RNA sequencing (scRNA-seq) analysis provided insights into intratumor and intertumor heterogeneity, highlighting the roles of specific cell types and intercellular communication patterns in resistance mechanisms. Moreover, a focal point of our study was the investigation of TFAP2B, one of the top genes identified in this model. We examined its role in modulating cisplatin resistance in ovarian cancer cells through in vitro experiments. Overexpression of TFAP2B in cisplatin-sensitive A2780 cells significantly enhanced their sensitivity to the drug, while overexpression in cisplatin-resistant A2780cis cells markedly inhibited cell growth. These findings underscore the potential of TFAP2B as a therapeutic target for overcoming platinum resistance in ovarian cancer.

## 2. Materials and Methods

The flow chart of the overall study was shown in Fig. S1. We elaborated on each step in the following sub-sections.

**Fig. S1.**
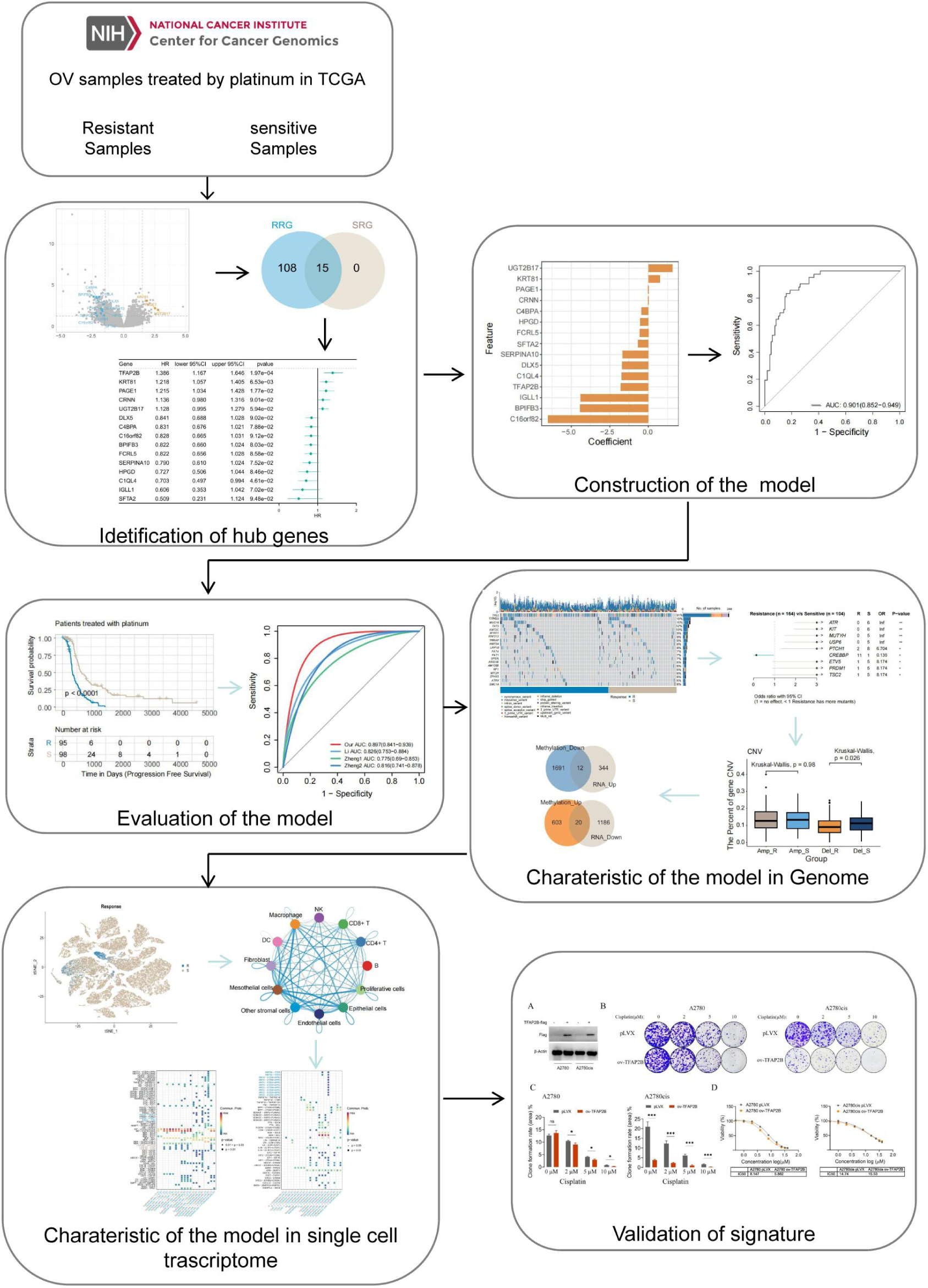
The workflow of this study.

### 2.1 Ovarian cancer Data Download and Preprocessing

Transcriptomic data were sourced from publicly accessible databases (Table 1), specifically The Cancer Genome Atlas (TCGA) and the Gene Expression Omnibus (GEO, transcriptome, methylation, copy number variation (CNV), and clinical data for TCGA-OV were retrieved from Xena (https://xenabrowser.net/datapages/). The selection process for the TCGA-OV dataset prioritized samples with complete survival records and extensive CNV, and transcriptomic information. Data on chemotherapy resistance, particularly to platinum-based regimens, were acquired from the cBioPortal for Cancer Genomics (https://www.cbioportal.org/), facilitating the analysis of genomic and epigenetic changes in relation to treatment efficacy and resistance mechanisms. For GEO datasets, probe-level expression data were consolidated to gene-level expressions using platform-specific annotations. In cases where multiple probes corresponded to a single gene, the expression values were averaged; instances of multiple genes mapping to a single probe were discarded. Further refinement was applied by filtering gene expression data through the HUGO Gene Nomenclature Committee database, ensuring the inclusion of only protein-coding genes. The TCGA-OV samples formed the basis of the training cohort for constructing the platinum resistance signature. Validation datasets comprised GSE17260, GSE30161, GSE18521, GSE32062, GSE26712, GSE63885, and GSE14764.

**Table 1.** The data sets of ovarian cancer used in this study.

| Data id | data type | Sample Status | OV samples | Data source |
| --- | --- | --- | --- | --- |
| TCGA-OV | RNA | Tumor (platinum Resistance/Sensitive) | 149 | TCGA |
| TCGA-OV | RNA | Tumor (survival) | 268 | TCGA |
| TCGA-OV | Methylation | Tumor | 7 | TCGA |
| TCGA-OV | CNV | Tumor | 268 | TCGA |
| GSE17260 | RNA | Tumor | 110 | GEO |
| GSE30161 | RNA | Tumor | 54 | GEO |
| GSE18521 | RNA | Tumor | 53 | GEO |
| GSE32062 | RNA | Tumor | 260 | GEO |
| GSE26712 | RNA | Tumor | 185 | GEO |
| GSE63885 | RNA | Tumor | 75 | GEO |
| GSE14764 | RNA | Tumor | 80 | GEO |
| scRNAseq-OV | scRNA | Tumor | 11 | Mendeley |

### 2.2 Chemotherapy Resistance-Related Driving Factors in Ovarian Cancer

The DESeq2 package in R was utilized for differential analysis on TCGA. With Benjamini-Hochberg (FDR) correction, genes with an FDR adjusted *P* value (adj.pvalue) < 0.05 and a log2 fold change (R vs S) of 1.5 were selected, identifying 123 chemotherapy resistance-related genes. The coxph function in the survival package was used to perform Cox proportional hazards analysis to calculate genes related to overall survival (OS) in TCGA-OV. Through the univariate Cox analysis, each target gene was evaluated as an independent prognostic factor in a regression analysis framework, facilitating the calculation of risk scores and the significance of each gene’s impact on survival outcomes. The analysis was carried out using the formula: coxph(formula = Surv(time, status) ∼ variable, data = clinical.data). A threshold of pvalue < 0.1 was used to select 15 survival-related genes from the resistance-associated genes. The ggVennDiagram package was employed to create a Venn diagram of differentially expressed genes and survival-related genes.

### 2.3 Construction of the Platinum Resistance Scoring Model and the Mutation and Epigenetic Characteristics based on its Groupings

To construction of the platinum resistance scoring model (PRSM), the glmnet package was used to build an elastic net model with 15 chemotherapy resistance-related genes, categorizing tumor patients in the training set cohort into sensitive and resistant groups. The predictive capability of the chemotherapy resistance scoring model was validated using ROC curves drawn by the pROC package. The model’s predictive efficacy was further validated using TCGA and GEO datasets, with survival curves plotted using the survminer package. Groupings were determined using the model’s optimal cutoff point for predicted scores. The TMB (Tumor Mutational Burden) score for each sample was first calculated using maftools. Then, using the maftools package, the mutation landscape of tumor driver genes in the chemotherapy resistance scoring model groups of the TCGA dataset was drawn, based on the tumor mutation gene list from the cosmic database CGC. Subsequent analysis involved conducting Fisher’s exact tests on the mutation information of the resistant and sensitive groups to identify differential mutation events. Data on HRD score and SNV_Neoantigen load differences were obtained from https://gdc.cancer.gov/about-data/publications/panimmune, and rank-sum tests were used to determine significant differences between resistant and sensitive groups. Analysis of somatic mutation types in typical cancer driver genes and CNV differences in the chemotherapy resistance scoring model groups followed. The ChAMP package was used to analyze methylation data. Differential methylation genes (with low methylation and high expression or high methylation and low expression) were selected for pathway enrichment analysis. The Δbeta threshold was set at 0.1, and the p-value had to be less than 0.05.

### 2.4 Survival analysis

Survival analysis was performed with the ‘survminer’ package in R, which facilitated the generation of survival curves using the Kaplan-Meier estimator for estimating non-parametric survival probabilities. This comprehensive package also enabled the utilization of the log-rank test, a statistical method for comparing survival curves between distinct groups delineated by clinical or molecular attributes. The integrated approach provided by ‘survminer’ allowed for a seamless analysis and visualization of survival data, reflecting the prognostic impact of various factors under investigation.”

### 2.5 Pathway Feature Analysis of Chemotherapy Resistance Scoring Model Classifications

The pathway feature analysis for the chemotherapy resistance scoring model was conducted based on the 50 hallmark pathways and GO biological process pathways (GOBP). Subsequently, the enrichment scores were calculated using the GSVA package, employing the ssGSEA method and assuming a poisson distribution for the data. Differences in enrichment scores were analyzed using the Wilcoxon rank-sum test. Following this, scores of HALLMARK gene sets exhibiting significant differences were subjected to Spearman’s rank correlation analysis alongside GOBP, to assess the strength and direction of association between them. The correlation network was visualized using CYTOSCAPE software (version 3.8.2, available at https://cytoscape.org/).

### 2.6 Single-cell data and preprocessing

Single-cell data for 11 ovarian tumor samples were obtained from the study by Zheng et al. [16]. Data preprocessing and analysis were conducted using the ’Seurat’ R package (version 4.0.4) as described by Hao et al. [17], applying default settings across all functions. Quality control measures involved the exclusion of cells with fewer than 200 detected genes or those with over 10% of unique molecular identifiers (UMIs) attributed to mitochondrial genes. The gene expression data of the filtered cells were subjected to log normalization and linear regression through the NormalizeData and ScaleData functions within the Seurat package (version 3.1.4). Cells exhibiting multiple major cell markers were considered doublets and were removed from the analysis on a cluster-by-cluster basis. The cells that passed these criteria were classified as single cells for further analysis. Cell type annotation was carried out based on established markers referenced in literature or sourced from the CellMarker and PanglaoDB databases.

### 2.7 Cell Communication Analysis

To dissect inter-cellular communication, our methodology incorporated the use of ’CellChat’ (version 1.5.0) [18]. After creating the CellChat object, Secreted signaling pathways were set as the reference database. We applied ’computeCommunProb’ to estimate cell-cell communication probabilities and strengths. Visualization of ligand-receptor interaction strengths was achieved through ’netVisual_bubble’.

### 2.8 Cell line and culture

Human ovarian cancer A2780 cell lines and A2780cis were cultured in complete medium consisting of DMEM, 10 % heat-inactivated FBS, 1% Pen/Strep. The cells were cultivated in an incubator with a temperature of 37*℃* and 95% air, with 5% CO2. Originally acquired from ATCC, the cells were routinely checked for mycoplasma infection. Short tandem repeat profiling was used to validate the cell lines, and mycoplasma contamination was examined. Cisplatin were purchased from Selleck Chemicals (Huston, TX, USA) and dissolved in dimethyl formamide (DMF).

### 2.9 Cell transfection and lentivirus infection

Lipofectamine 3000 (Gibco, Invitrogen) was used for the transfection experiment according to the manufacturer’s instructions. TFAP2B was generated by PCR amplification and cloned into the pLVX-3×FLAG-Puro vector for overexpression with the following primers: hTFAP2B_rec forward primer (5′-GTGAATTCCTCGAGAGCCACCATGCACTCACCTCCTAGAGACC-3′) and hTFAP2B_rec reverse primer (5′-ACGCGTGTATACGTTTCATTTCCTGTGTTTCTCCTCCTTG-3′) . The 293 T cells were transfected with pLVX-TFAP2B-3×Flag together with psPAX2 and pMD2.G lentiviral packaging systems (Addgene) to generate TFAP2B stably overexpressed cell lines.

### 3.0 Western Blot

For protein quantification, cells were lysed in RIPA buffer (50 mmol/L Tris-HCI, 150 mmol/L NaCl, 0.5 mmol/L EDTA, 1% NP40, 10% glycerin), and Pierce, Waltham’s BCA Protein Assay Kit was utilized. Proteins of the same amount were added to the loading buffer, brought to a boil at 95°C for 10 minutes, separated using SDS-PAGE, and then put onto a PVDF membrane (Bio-Rad, Hercules). The membrane was blocked for one hour at room temperature with 5% defatted milk, and then it was incubated with primary antibodies, such as β-actin (Cell Signaling Technology, Boston) and flag (Sigma, USA), for the entire night at 4°C. Secondary antibodies (Bio-world, Minnesota) were incubated for an additional hour at room temperature following three rounds of TBST washing. Luminescence Imaging System (Tanon, Shanghai) was used to identify target protein bands.

### 3.1 CCK8 and Crystal violet cell viability and proliferation assay

Each cell line was seeded at a density of 4–6×10^2^ cells/well in flat-bottomed 96 well culture plate in 100uL of the culture medium. Stock solutions of cisplatin were subjected to serial dilutions to give final concentrations ranging from 0 μM to 40 μM. The plate was further incubated for 72 hours in the CO_2_ incubator. Subsequently, 10 μL of CCK-8 solution was added to each well, ensuring no bubble formation to avoid interference with optical density (OD) readings. The plate was then returned to the CO_2_ incubator and incubated for an additional 4 hours. Absorbance at 450 nm was measured using a microplate reader to assess cell viability. For the colony-formation assay, cells were seeded in 6-well plates (200 cells/well) and treated with different concentration of cisplatin ranging from xx to xx for 10 to 14 days depending on growth rate. After 14 days, cells were fixed with methanol and stained with 0.1% crystal violet. The area of colonies was then quantified for analysis.

### 3.2 Statistical analysis

Statistical analyses were executed using R software (versions 4.3.1). Differences in clinical characteristics between the training and internal validation cohorts were assessed using the chi-squared test. For comparing two groups with non-normally distributed variables, the Wilcoxon test served as the non-parametric method of choice. Differentially expressed genes (DEGs) were identified using an FDR-corrected p-value to determine statistical significance. Kaplan–Meier survival curves and log-rank tests, facilitated by the R “survival” package, were employed to evaluate overall survival (OS) across various subgroups. Univariate Cox regression analysis was applied to discern independent prognostic indicators. The efficacy of the predictive model was appraised through ROC curve analysis and area under the curve (AUC) computation, utilizing the “timeROC” R package. Spearman’s correlation analysis was conducted to elucidate the relationships between hallmark and GOBP pathways. Student’s t-test was used in colonies formation rate ananlysis, a p-value of <0.05 was predetermined as the threshold for statistical significance, unless specified otherwise.

## 3. Results

### 3.1 DEG and prognosis-based platinum signature for ovarian cancer

To elucidate the genetic factors contributing to platinum resistance in ovarian cancer, we performed a differential expression genes (DEGs) analysis comparing platinum-resistant and platinum-sensitive groups within the TCGA-OV database. This analysis included 107 patients who exhibited a positive response to chemotherapy, categorized as the platinum-sensitive group (S), and 42 patients who did not respond to chemotherapy, defined as the platinum-resistant group (R). We identified differentially expressed genes by applying a threshold of an adjusted p-value (Benjamini-Hochberg FDR) of <0.05 and a |log2 fold change (R vs S)| of 1.5. This approach yielded 123 differential expressing genes associated with chemotherapy resistance (Fig. 1A), which were subsequently considered for inclusion in a platinum-resistance gene panel. In an effort to discern the key genes impacting the prognosis of ovarian cancer patients undergoing chemotherapy, we evaluated 268 patients from the TCGA-OV database. Using these data, we developed univariate Cox proportional hazard regression models (Supplementary Table S1). Of the 123 chemotherapy resistance-related genes previously identified, 15 were found to be significantly associated with overall survival (OS) according to the univariate Cox regression analysis results (Fig 1B, C). Among the 15 genes, KRT81, PAGE1 and UGT2B17 were up-regulated in platinum-resistant group (orange), while the remaining ones were down-regulated (blue) as shown in Fig.1A. Then, as shown in the forest plot of the 15 platinum resistance-related genes, hazard ratios exceeding 1 suggest that patients exhibiting upregulation or downregulation of these genes (TFAP2B, KRT81, PAGE1, CRNN, UGT2B17) are at an increased risk of tumor progression after receiving platinum-based chemotherapy (Fig. 1C).

**Fig. 1:**
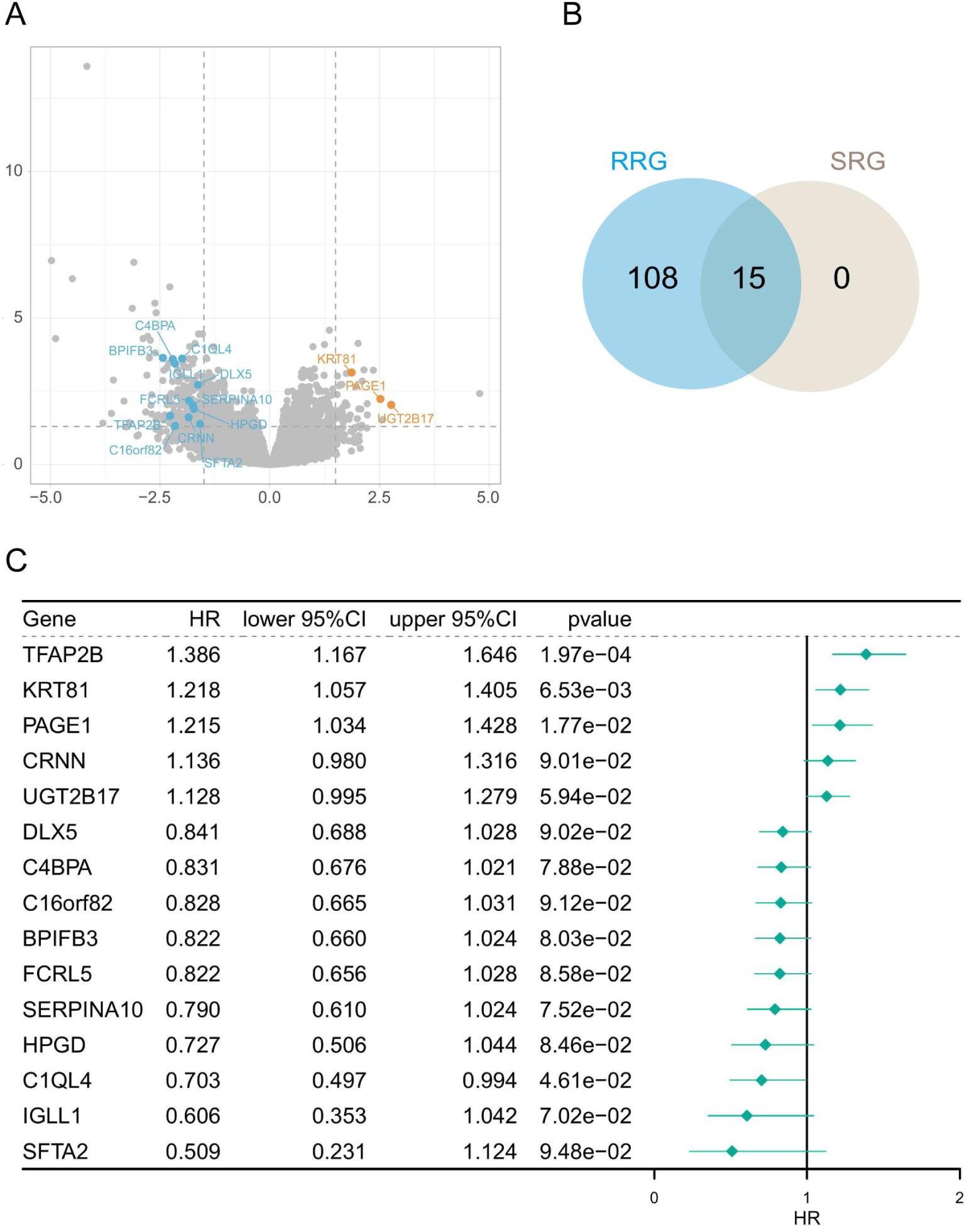
The genomic landscape distinguishing platinum-resistant from platinum-sensitive cases in ovarian cancer. (A) Volcano plot showcasing differential gene expression between the platinum-resistant group and the platinum-sensitive group in TCGA, based on adjusted adj.pvalue (≤0.05) and |Log2 fold change| (≥ 1.5). (B) Venn diagram depicting the overlap between differentially expressed genes and genes associated with overall survival, identifying 15 key genes implicated in both platinum resistance and patient survival outcomes. (C) Forest plot of the 15 platinum resistance-related genes, demonstrating their hazard ratios and 95% confidence intervals, indicating the strength and direction of each gene’s association with survival in patients undergoing chemotherapy.

### 3.2 Analysis of ceRNA networks and TF regulatory networks associated with 15 platinum resistance-related genes

In our comprehensive investigation, we delved into the ceRNA networks and transcription factor (TF) regulatory frameworks related to fifteen pivotal hub genes. The initial step involved sourcing tumor-associated microRNAs (miRNAs) from the Human MicroRNA Disease Database (HMDD), leading to the extraction of 1,273 mRNA-miRNA pairs associated with the hub genes’ mRNA, as cataloged in the miRWalk database. A subsequent intersection with 364 tumor-associated miRNAs yielded 7 significant mRNA-miRNA pairs (depicted in Fig. S2A). Notably, the implicated miRNAs predominantly belong to the hsa-let-7 family, including hsa-let-7a, hsa-let-7b, hsa-let-7c, hsa-let-7d, hsa-let-7e, hsa-let-7g, and hsa-let-7i. This finding aligns with existing research that highlights the critical role of the human let-7 family - comprising 13 members spread across nine different chromosomes - in modulating drug sensitivity and resistance, particularly in relation to platinum-based chemotherapy. For instance, deregulation of let-7e in epithelial ovarian cancer has been implicated in the development of cisplatin resistance, underscoring the intricate relationship between miRNA expression and chemotherapy efficacy [19]. Further investigations, utilizing tumor-associated miRNAs and the ENCORI database, identified interacting long non-coding RNAs (lncRNAs), culminating in the delineation of lncRNA-miRNA-mRNA interaction networks (illustrated in Fig. S2B, Supplmentary table S2). Moreover, motif enrichment analyses revealed that the expression of hub genes might be under the regulatory influence of transcription factors such as FOXII, SOX4, and ZNF234 (Fig. S2C). Intriguingly, SOX4 has been identified as a potential therapeutic target in cervical cancer, implicated in modulating cancer cell sensitivity to drugs through the regulation of ABCG2 [20]. Similarly, ZNF234, known for its roles in DNA binding and transcriptional regulation, has been associated with cellular responses to chemical stimuli, further emphasizing the complexity of gene regulation in the context of cancer therapy resistance [21].

### 3.3 Platinum-resistant samples classified by the Platinum Resistance Scoring Model (PRSM) showed worse prognosis

Utilizing an elastic net regression approach, we developed a novel platinum resistance scoring model to enhance the prediction of treatment responses in ovarian cancer. This model was finely tuned to optimal performance with an alpha of 0.2 and lambda set to zero, enabling precise determination of the influence of 15 hub genes on chemotherapy resistance (Fig. 2A). The calculated coefficients for these genes provide a quantitative measure of their respective contributions to the model’s predictive capability, thereby establishing a robust framework for resistance assessment. The validity of the model was assessed through the construction of receiver operating characteristic (ROC) curves, resulting in an area under the curve (AUC) of 0.901 (Fig. 2B). Extensive validation of the model’s performance was conducted by comparing the predicted resistance scores between patient groups categorized as sensitive or resistant based on their treatment outcomes. The scoring model effectively segregated these groups with a wilcoxon rank-sum test’s P value less than 0.05, demonstrating its capability to accurately reflect the underlying genetic determinants of chemotherapy response (Fig. 2C). Further validation was conducted within the TCGA dataset, comprising both a training set with documented drug sensitivity and the comprehensive patient cohort. The model proficiently stratified patients, demonstrating its utility in predicting clinical outcomes based on overall survival (OS) and progression-free survival (PFS), with significant distinctions between patient prognoses (Fig 2D-F).

**Fig. 2:**
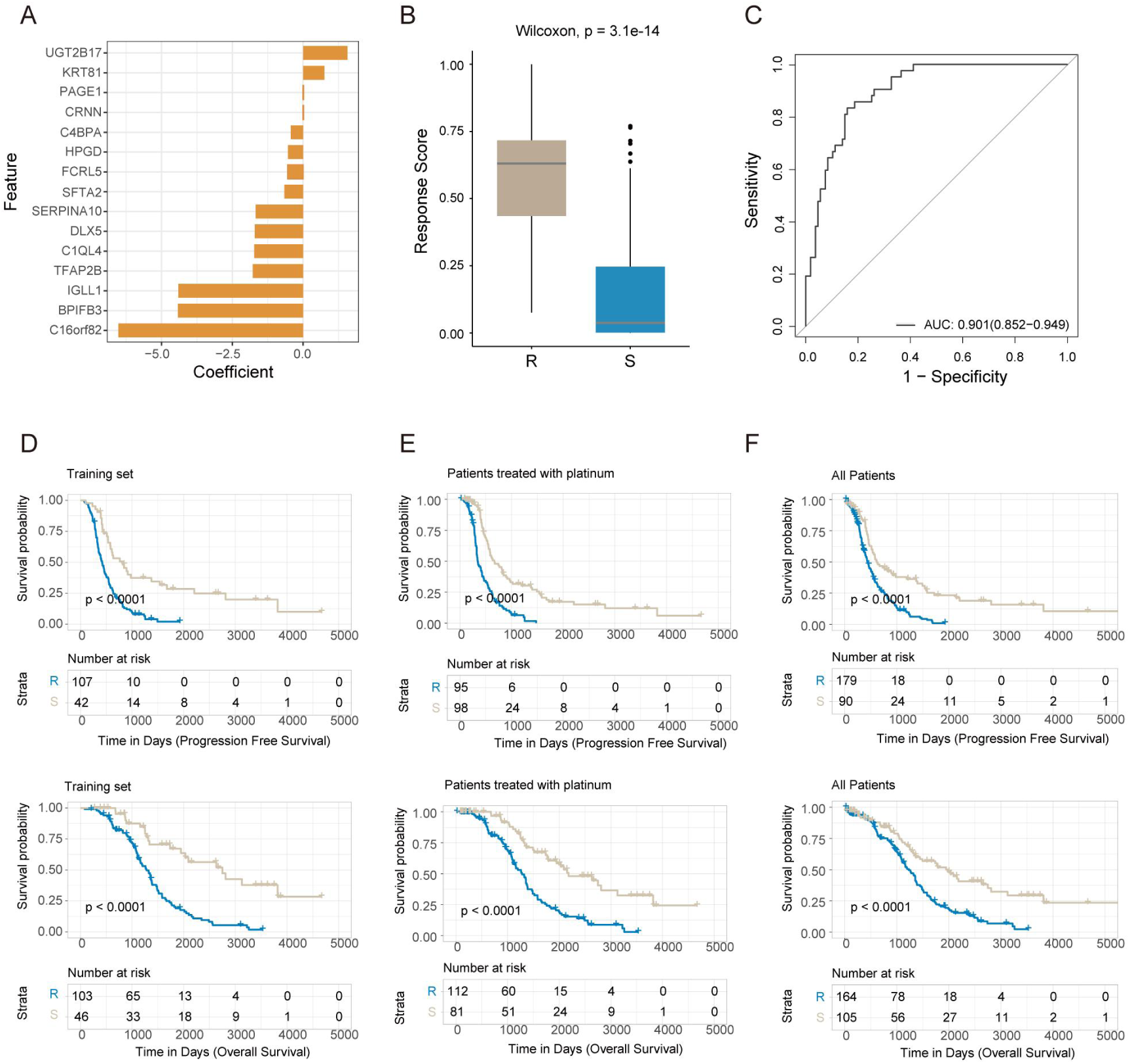
Construction and validation of the Platinum Chemotherapy Resistance Scoring Model in OV patients (A) Coefficients of 15 hub genes within the elastic net regression model predicting platinum chemotherapy resistance. (B) Distribution of predicted resistance scores in the TCGA dataset, comparing patients classified into platinum-resistant and platinum-sensitive groups based on the model’s scoring (Wilcoxon rank sum test, *P* = 3.1e-14). (C) The ROCs for the TCGA training cohort (149 samples), demonstrating the elastic net regression model predictive accuracy with an area under the curve (AUC = 0.901) metric. ROC, receiver operating characteristic; TCGA, the cancer genome atlas. (D-F) Kaplan-Meier survival curves for patients stratified by the model into resistant and sensitive groups, showing differences in progression-free survival (PFS) and overall survival (OS) across the groups classified by 15 hub genes in TCGA training data, TCGA platinum-treated data and TCGA all patients data. R, resistant samples; S, sensitive samples.

### 3.4 PRSM signature was futher validated and showed better performance than other platinum signatures

Moreover, PRSM’s efficacy was further corroborated through analysis across seven independent OV datasets, totaling 817 patients from the GEO, where it consistently delineated patient groups with distinct prognoses based on the Platinum Resistance Scoring Model (PRSM) (GSE17260, *P* = 0.0018; GSE14764, *P* = 0.047; GSE30161, *P* = 0.042; GSE26712, *P* = 0.0033; GSE18521, *P* = 0.094; GES32062, *P* = 0.021; GSE63885, *P* = 0.033. Fig. 3A). Besides, a comparative assessment with previously established signatures—specifically those by Zheng et al. [22] and Li et al. [23]—revealed PRSM model’s superior predictive efficiency. In TCGA dataset, PRSM model’s AUC was 0.897 and in GSE63885 data set AUC was 0.626 (Fig 3B-C). Such comparisons, spanning multiple datasets and existing models, affirm the robustness and enhanced predictive capability of our platinum resistance scoring model, highlighting its potential to refine therapeutic strategies for ovarian cancer.

**Fig. 3:**
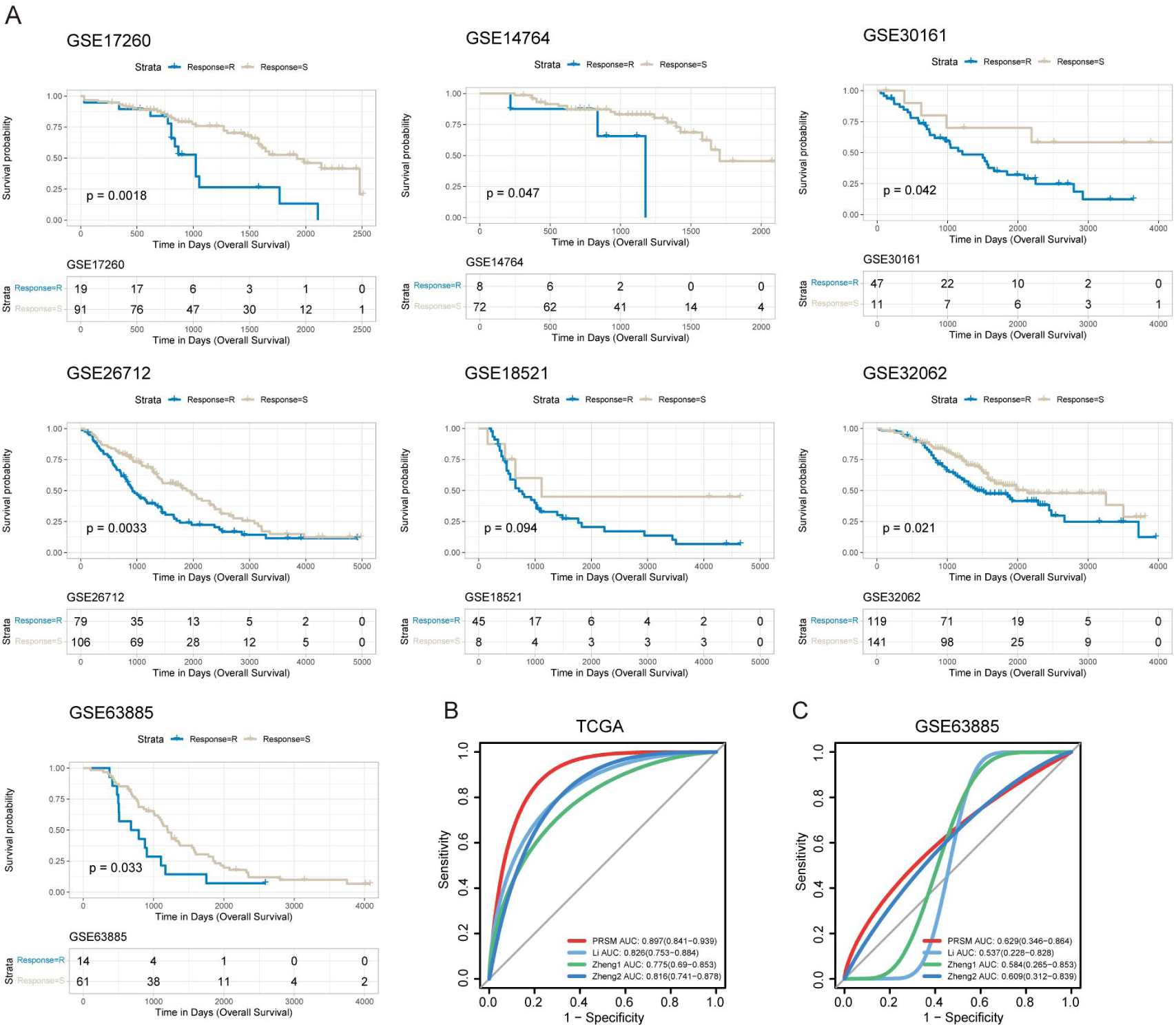
Extending the analysis of the chemotherapy resistance scoring model by demonstrating its applicability and performance beyond the initial dataset. (A) Validation of the model using Gene Expression Omnibus (GEO) datasets, showcasing the predictive capability of the model in identifying platinum-resistant and platinum-sensitive cases in external patient cohorts. Kaplan–Meier survival curves show the OS difference between platinum-resistant and -sensitive OV samples classified by 15 hub genes in GSE17260, GSE30161, GSE18521, GSE32062, GSE26712, GSE63885, and GSE14764. (B-C) Comparative analysis of the PRSM model’s performance against other published signatures within the TCGA (149 samples) and GSE63885 dataset (75 samples). PRSM signature (red).

### 3.5 Platinum-resistant samples classified by PRSM exhibited lower Tumor Mutational Burden (TMB) and lower Single Nucleotide Variant (SNV) neoantigen loads

In this section, the focus was on the mutation and epigenetic characteristics of the groups classified by the platinum resistance scoring model. The mutation landscape of the top 20 most frequently mutated tumor driver genes, selected from a list of 738 genes in the cosmic database CGC, was depicted for the TCGA dataset groups using maftools (Fig. 4A). Fisher’s exact test identified significant disparities in the mutation rates of 30 genes, such as ATR, KIT, MUTYH, USP6, PTCH1, CREBBP, ETV5, PRDM1, and TSC2, across the groups (*P* < 0.05, Fisher’s exact test, Fig. 4B). Among these genes, CREBBP has more mutants in the resistant group with P-value less than 0.05 (Fig. 4B). Additionally, copy number variation (CNV) analysis showed the frequencies of deletion in the resistant samples were significantly lower than those in the sensitive samples (*P* = 0.026, Kruskal–Wallis test, Fig. 4C), while frequencies of amplification showed no significantly difference (*P* = 0.98, Kruskal–Wallis test, Fig. 4C). Further analyses revealed that, compared to the sensitive group, the resistant group exhibited a significantly lower Tumor Mutational Burden (TMB) (*P* = 0.044, Wilcoxon rank-sum test, Fig. 4D). Additionally, the resistant group had lower SNV Neoantigen load (*P* = 0.096, Wilcoxon rank-sum test, Fig. 4E) and a lower Homologous Recombination Deficiency (HRD) scores (*P* = 0.42, Wilcoxon rank-sum test, Fig. 4F), although these differences were not statistically significant.. Besides, by comparing the methylation profiles between platinum-resistant and -sensitive OV samples classified by PRSM in TCGA, 623 hypermethylated and 1703 hypomethylated genes were identified respectively, based on ΔBeta values (>0.1 or <-0.1) and *P*-values (<0.05) (Fig. 4G). Subsequently, RNA expression analysis showed 356 upregulated and 1206 downregulated genes. Integrating methylation and RNA expression results led to the identification of genes with low methylation and high expression (12 genes), and high methylation with low expression (20 genes), followed by pathway enrichment analysis (Fig. 4G). The enrichment analysis indicated that genes upregulated in the resistant group were primarily involved in substance transport processes (Fig. 4H), while downregulated genes were mainly associated with morphogenesis (Fig. 4I).

**Fig. 4:**
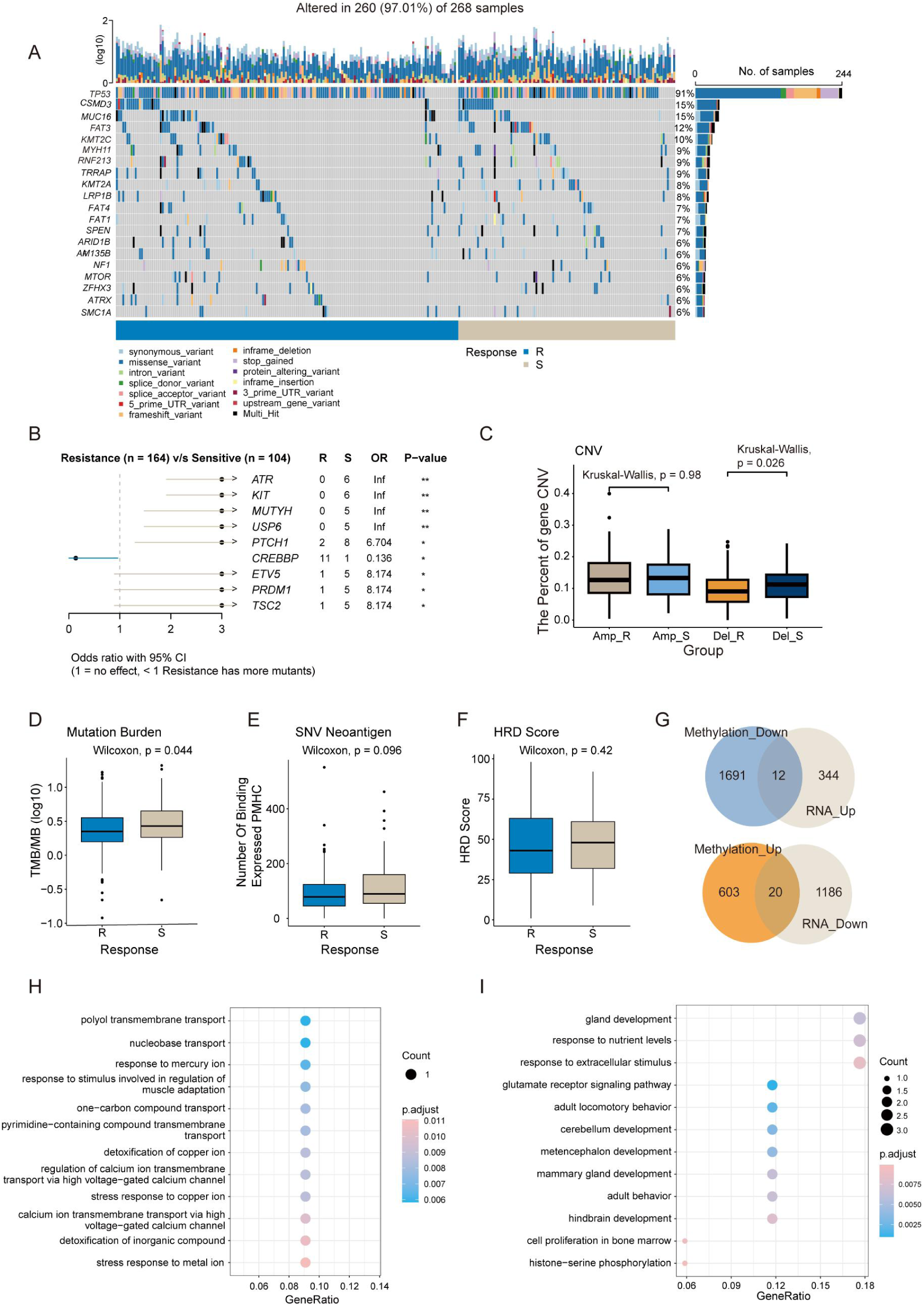
The multiomics landscape between platinum-resistant and -sensitive OV samples classified by PRSM in TCGA. OV, ovarian cancer; PRSM, 15 genes based platinum chemotherapy resistance scoring model ; TCGA, the cancer genome atlas. (A) A mutation waterfall plot for the top 20 tumor driver genes shows the mutation frequency and distribution across samples in the model-defined groups, with genes as rows and samples as columns. The mutation percentage difference between platinum-resistant (R) and -sensitive (S) samples is depicted in the right bar, while the total mutation count is shown in the top bar. R, resistant samples; S, sensitive samples. (B) A forest plot presents the hazard ratios and confidence intervals for mutated genes, indicating their potential influence on chemotherapy resistance. R, resistant samples; S, sensitive samples; Fisher’s exact test, \**P* < 0.05, \*\**P* < 0.01; OR, odds ratio with 95% CI (1 = no effect, < 1 Resistance has more mutants); (C) Differences in copy number variation (CNV) amplification and deletion between platinum-resistant and sensitive samples in the TCGA, with the median, quartiles, and range depicted. The vertical line reaches the maximum and minimum values. CNV, copy number variation; Amp_R, amplification in resistant samples; Amp_S, amplification in sensitive samples; Del_R, deletion in resistant samples; Del_S, deletion in sensitive samples; (D) Tumor Mutational Burden (TMB) comparison between platinum-resistant and sensitive samples in the TCGA (Wilcoxon rank sum test, *P* = 0.044). (E) Comparison of Single Nucleotide Variant (SNV) neoantigen counts between resistant and sensitive groups in the TCGA (Wilcoxon rank sum test, P=0.096). (F) Distribution of Homologous Recombination Deficiency (HRD) scores across platinum-resistant and sensitive groups in the TCGA (Wilcoxon rank sum test, *P* = 0.42). (G) A Venn diagram showing the overlap of differentially methylated positions and differentially expressed genes between resistant and sensitive groups. (H-I) Pathway enrichment diagrams for genes that are respectively upregulated and downregulated, highlighting the biological processes and pathways preferentially activated or suppressed in resistant versus sensitive groups.

### 3.6 Platinum-Resistant Samples Classified by PRSM showed Metabolic Pathway Alterations

Our investigation extended to analyzing pathway features for the chemotherapy resistance scoring model from the 50 Hallmark gene sets. Through single-sample gene set enrichment analysis (ssGSEA), distinct pathways were identified as being significantly enriched in the platinum-resistant group compared to the sensitive group. These pathways included MYC targets V2 and Oxidative Phosphorylation in resistant group, while Hypoxia, Notch signaling Apical surface, Hedgehog signaling, Fatty acid metabolsim, Xenobiotic metabolism, Spermatogensis, Kras signaling DN in sensitive group (*P* < 0.05, Wilcoxon rank-sum test, Fig. 5A). Furthermore, we performed correlation analyses between these 11 pathways and Gene Ontology Biological Processes (GOBP) within both the resistant and sensitive groups. In the resistant cohort, we observed a notable presence of positive correlations with pathways such as the pyrimidine nucleobase catabolic process, intrinsic apoptotic signaling pathway, and vascular-associated smooth muscle cell apoptotic process—correlations that were conspicuously absent in the sensitive group (Fig. 5B and 5C, Supplmentary table S3). Conversely, pathways related to metabolism, including rRNA base methylation, cardiolipin biosynthetic process, and flavone and flavonoid metabolic processes, exhibited negative correlations within the resistant group (Fig. 5B). This delineation underscores a distinct biochemical landscape between the resistant and sensitive groups, emphasizing the complex interplay of cellular processes in mediating platinum chemotherapy resistance.

**Fig. 5:**
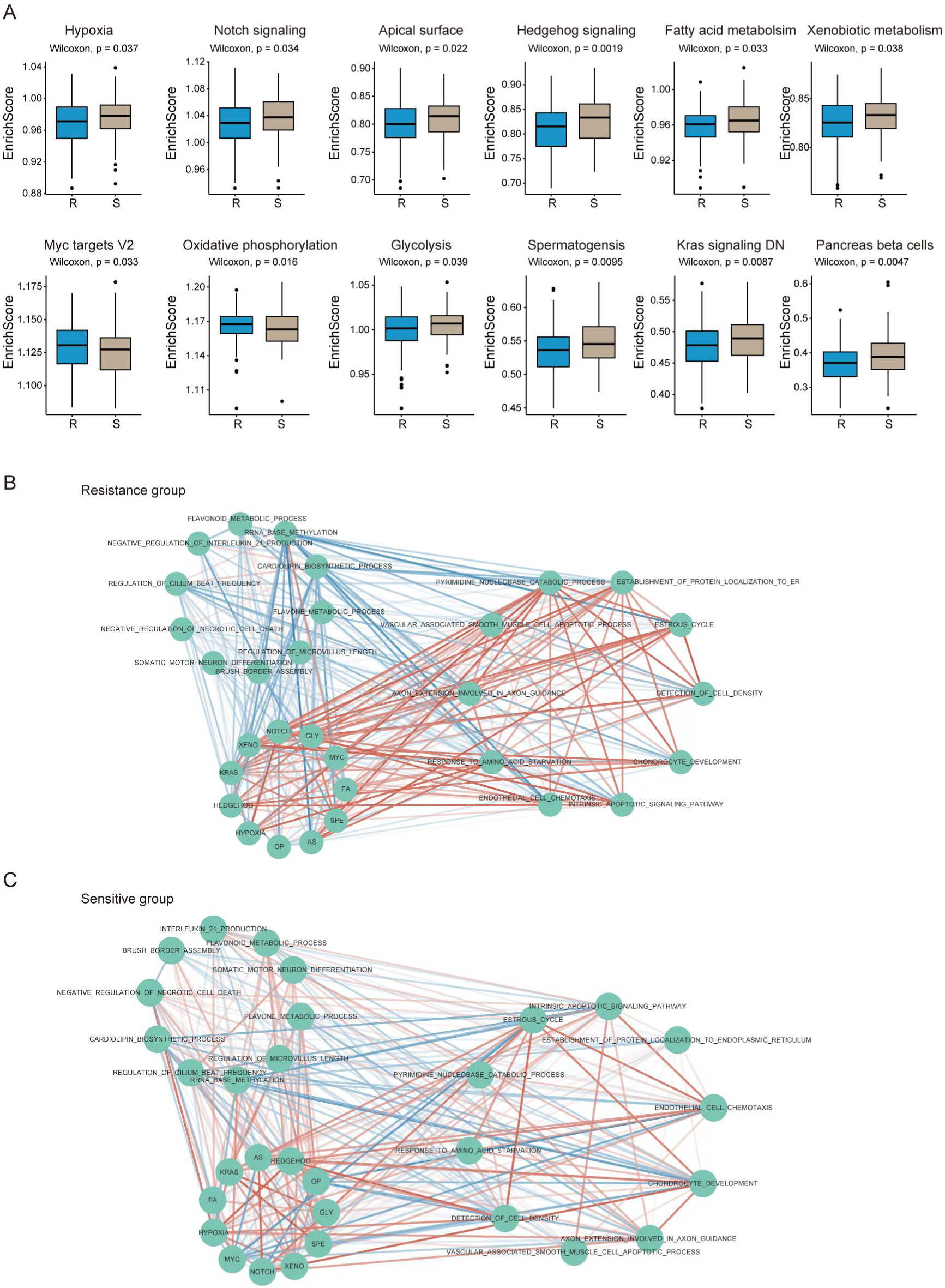
Pathway Feature Analysis of Chemotherapy Resistance Scoring Model Classifications in Ovarian Cancer. This figure provides an in-depth pathway analysis distinguishing platinum-resistant from platinum-sensitive groups, based on our chemotherapy resistance scoring model. (A) Differential pathway enrichment scores between resistant and sensitive groups were determined using single-sample Gene Set Enrichment Analysis (ssGSEA) based on hallmark gene sets from The Molecular Signatures Database (MSigDB). R, resistant samples; S, sensitive samples. (B) Network diagram illustrating the correlation between differentially enriched pathways in the resistant group and Gene Ontology Biological Processes (GOBP), highlighting the complex interplay of pathways that contribute to chemotherapy resistance. (C) Corresponding network diagram for the sensitive group, demonstrating the distinct pathway and Gene Ontology Biological Processes (GOBP) associations unique to this classification.

### 3.7 Single-Cell Analysis of Intratumor and Intertumor Platinum Resistance Heterogeneity

Considering the notable disparities between platinum-resistant and sensitive ovarian cancer (OV) samples, our objective was to explore how cellular diversity influences platinum resistance on a single-cell level. After quality control, we retained a total of 48,881 cells from primary OV tumor samples, which were annotated into 14 clusters (Fig. 6A). We observed substantial heterogeneity in the cellular composition across different samples (Fig. 6B). The hub genes associated with platinum based chemotherapy resistance in ovarian cancer were primarily expressed in epithelial cells (cancer cells), NK cells, and other cell types (Fig. 6C). However, the expression level of TFAP2B was too low to be detected in these single-cell transcriptomes. Subsequently, utilizing 15 pivotal genes, we classified cells into two subtypes, with NK and epithelial cells showing a pronounced enrichment in the resistant subgroup (Fig. 6D). This enrichment suggests a crucial role for these cell types in mediating chemotherapy resistance in OV. Furthermore, cells within the same sample exhibited traits of both sensitivity and resistance to platinum (Fig. 6E), echoing the findings of a recent study by Wang et al. [24], thereby highlighting the complex interplay of cellular characteristics in response to chemotherapy.

**Fig. 6:**
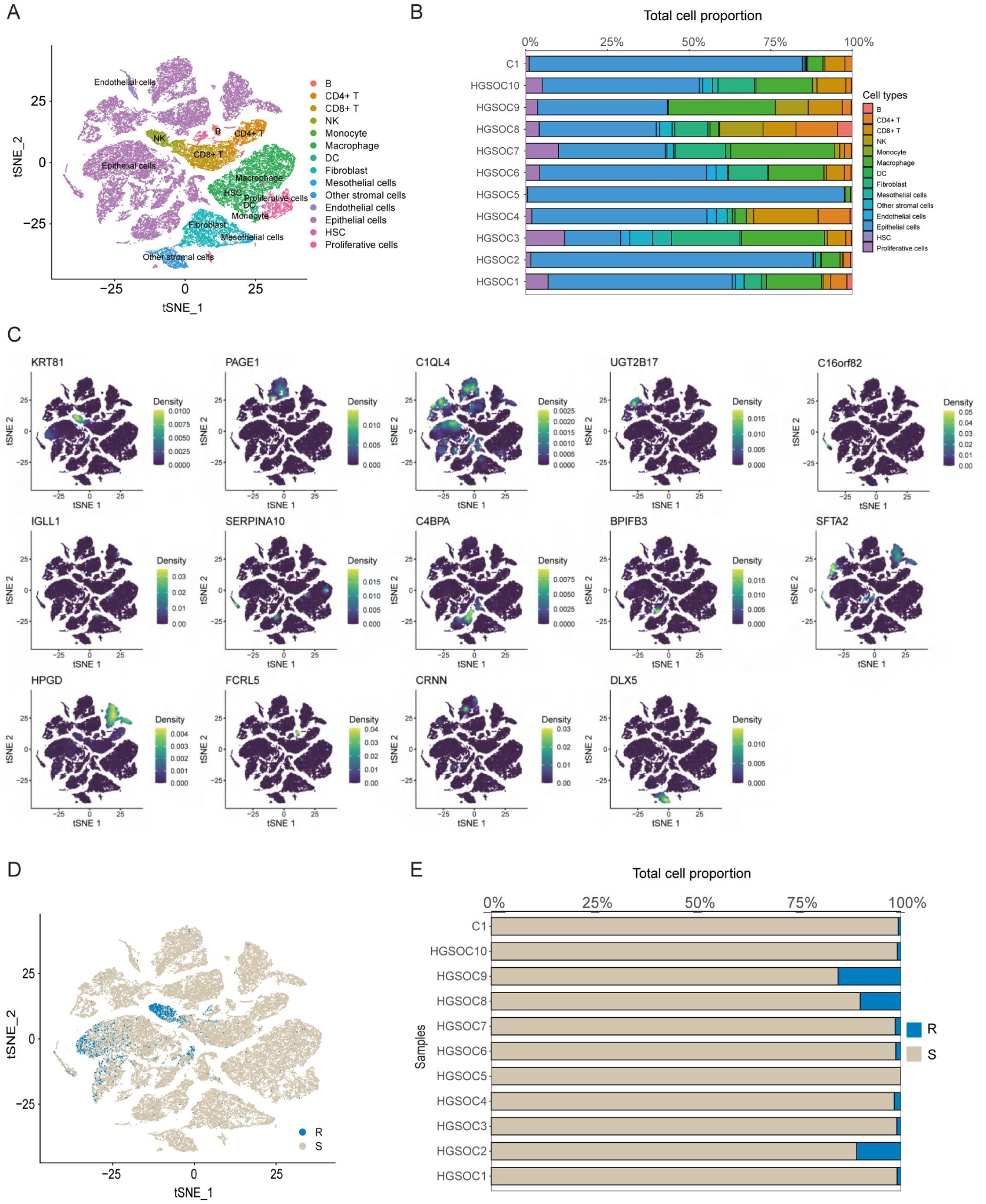
Single-Cell Analysis of Intra- and Intertumoral Heterogeneity in Platinum Chemotherapy Resistance. This figure explores the cellular heterogeneity of platinum chemotherapy resistance within and across ovarian cancer tumors through single-cell RNA sequencing (scRNA-seq) analysis. (A) t-Distributed Stochastic Neighbor Embedding (t-SNE) visualization of cells analyzed by scRNA-seq, categorizing them into distinct clusters to illustrate the diversity in cell types present within ovarian cancer samples. (B) Bar graph showing the relative proportions of cell types identified across 11 ovarian cancer (OV) samples, highlighting the variability in cellular composition among tumors. (C) t-SNE plot depicting the expression patterns of 14 hub genes associated with chemotherapy resistance across different cell types. The size and color of the bubbles indicate the level of gene expression and the prevalence of each gene within specific cell types, respectively. (D) t-SNE visualization of cells classified according to the chemotherapy resistance predictive model, demonstrating the spatial distribution of resistant versus sensitive cells within the t-SNE plot. (E) Bar graph presenting the relative proportions of resistant and sensitive cells within the 11 OV samples analyzed, offering insights into the extent of heterogeneity in chemotherapy response at the single-cell level.

### 3.8 Cell Communication Analysis Reveals Mechanisms of Chemotherapy Resistance

Beyond intrinsic cellular characteristics, cell-cell interactions may also influence platinum resistance. Our analysis revealed a marked increase in communication between epithelial cells and both mesothelial and immune-related cells within the resistant group, a phenomenon not observed in the sensitive cohort (Fig. 7A). Next, by comparing the interaction strengths of each signaling pathway, specific environmental signaling pathways between the resistant and sensitive groups were identified. In the resistant group, a broad activation of signaling pathways was noted, including but not limited to LC, ncWNT, CD70, TWEAK, and WNT, with a notable emphasis on the CCL signaling pathway being predominantly active in the sensitive group (Fig. 7B). Conversely, the majority of other signaling pathways remained inactive in the sensitive cohort. Delving deeper into the receptor-ligand interactions between cancer cells and the tumor microenvironment, it was revealed that the PTN-NCL axis was specifically engaged on cancer cells within the sensitive group (Fig. 7C). Additionally, within the context of WNT signaling, ligands such as WNT5A, WNT4, and WNT2, in conjunction with their receptors FZD6/FZD3, were particularly active in bridging mesothelial to epithelial cell communication in the resistant group. On the contrary, the resistant group also displayed activation of the PRSS1 ligand and receptors including F2R, F2RL1, and PARD3, facilitating signaling from epithelial to mesothelial cells (Fig. 7C-D). Meanwhile, the LGALS9 ligand alongside its receptors HAVCR2/CD44/CD45 exhibited significant activity in the sensitive group, enhancing communication from epithelial cells to various immune cells such as B cells, CD4+ T cells, CD8+ T cells, NK cells, macrophages, and dendritic cells (Fig. 7D). This nuanced exploration highlights the intricate signaling interplays that define the resistant versus sensitive group dynamics, offering insights into the molecular underpinnings of chemotherapy response.

**Fig. 7:**
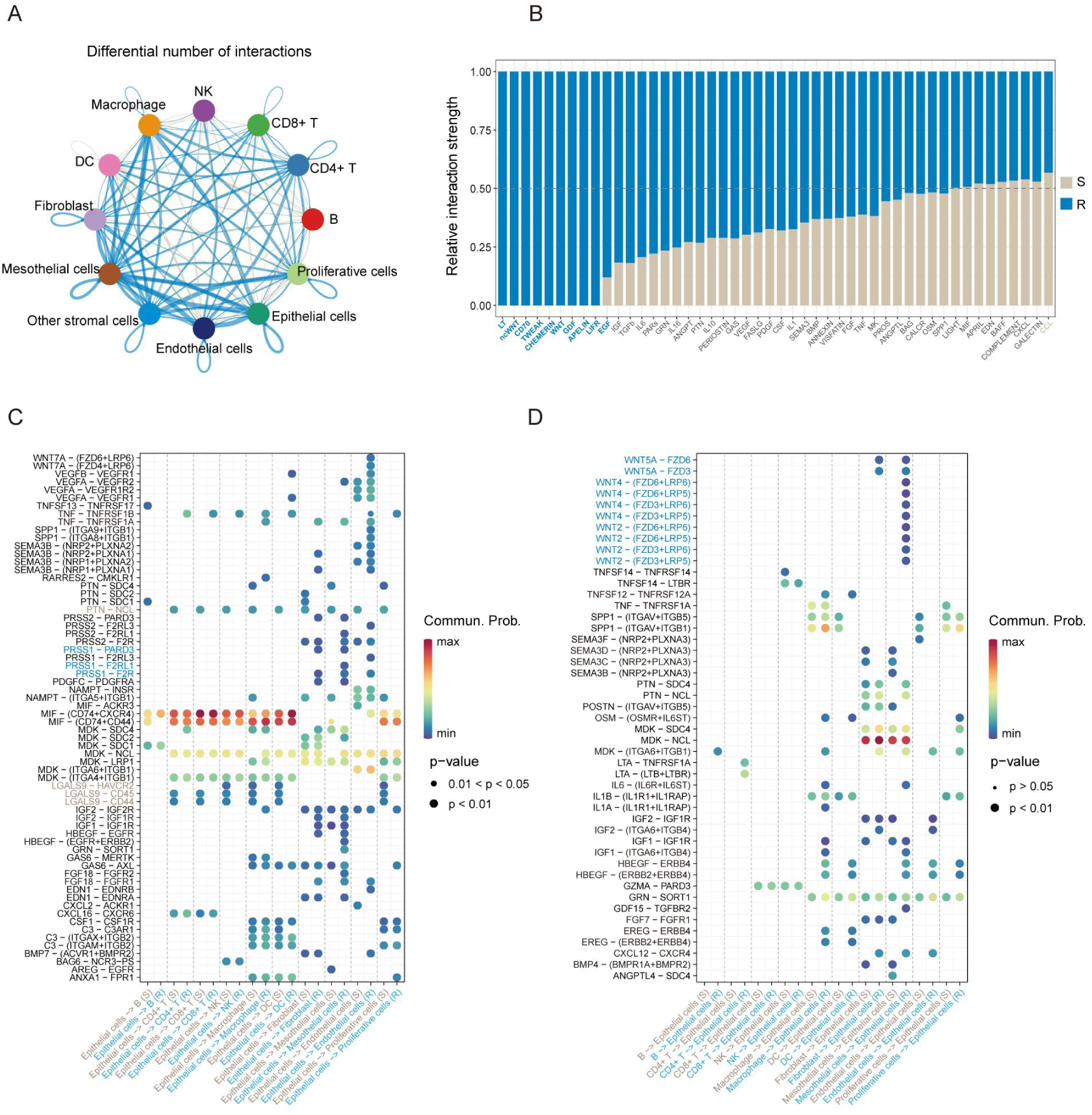
Cell Communication Analysis Reveals Mechanisms of Chemotherapy Resistance. This figure investigates the differential cell-cell communication patterns between platinum-resistant and -sensitive groups in ovarian cancer, elucidating potential mechanisms underlying chemotherapy resistance. (A) Overview of ligand-receptor interactions comparing the resistant group to the sensitive group. Interactions are visualized with lines, where blue indicates a higher number of interactions in the resistant group compared to the sensitive group, and light brown represents the opposite. The thickness of each line correlates with the difference in interaction count. (B) Bar graph depicting the relative interaction strengths of various signaling pathways between the chemotherapy-resistant and -sensitive groups. Pathways enriched in the resistant group are shown in blue at the top, while those enriched in the sensitive group are in light brown. (C and D) Comparison of ligand-receptor pairs between cancer cells and the tumor microenvironment (TME) cells in the resistant and sensitive groups, illustrating the dynamics of intercellular communication. (C) Focuses on communication from cancer cells to TME cells, while (D) highlights communication from TME cells to cancer cells. The significance of each interaction is represented by the size of the circles (P-values), and the probability of communication is indicated by the color of the circles.

### 3.9 Enhanced Sensitivity to Cisplatin Resistance in Ovarian Cancer Cells Overexpressing TFAP2B from PRSM signature

Among the 15 core genes identified in Fig. 1C, associated with drug resistance and prognosis, five genes (TFAP2B, KRT81, PAGE1, CRNN, UGT2B17) were found to be linked to poor prognosis in platinum-resistant ovarian cancer. Notably, among the 5 genes, TFAP2B exhibited the highest contribution to the Platinum Resistance Scoring Model (PRSM) constructed in Fig. 2A, as evidenced by its highest coefficient absolute value. Consequently, TFAP2B was selected for molecular biology validation to ascertain its association with platinum resistance in ovarian cancer. To elucidate the role of TFAP2B in cisplatin resistance, we over-expressed TFAP2B in cisplatin sensitive A2780 and cisplatin resistant A2780cis cell lines (Fig. 8A, Supplementary file 1). Subsequently, colony formation assays revealed that TFAP2B overexpression enhanced the responsiveness of A2780 ovarian cancer cells to cisplatin treatment (Fig. 8B, C). Meanwhile, we examined the effect of TFAP2B overexpression in A2780cis cisplatin-resistant cells. Remarkably, TFAP2B overexpression significantly inhibited proliferation of A2780cis cells as observed in colony formation assays (Fig. 8B, C). Co-treatment with cisplatin further suppressed the growth of A2780cis cells. Besides, TFAP2B overexpression in A2780 cells led to a significant reduction in cisplatin IC50 values in TFAP2B-overexpressing A2780 cells (Fig. 8D), indicating increased sensitivity to cisplatin. Notably, overexpression of TFAP2B in A2780cis cells did not significantly alter the IC50 value for cisplatin resistance (Fig. 8D). These findings corroborate the significance of TFAP2B in modulating cisplatin resistance in ovarian cancer and underscore its potential as a therapeutic target for overcoming drug resistance in this malignancy.

**Fig. 8:**
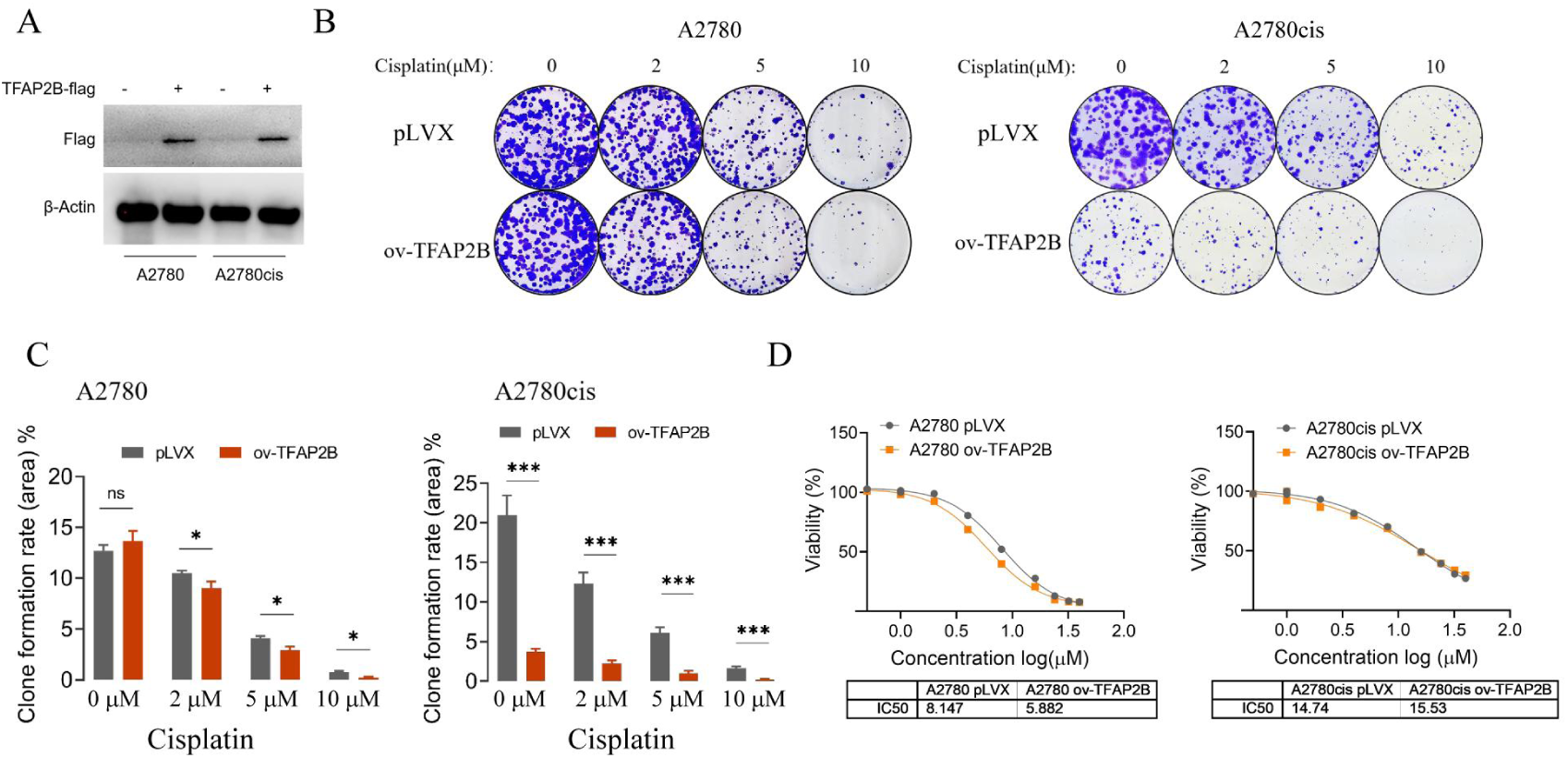
Cisplatin-sensitivity of TFAP2B knockin A2780 and A2780cis cell lines. (A) Verification of TFAP2B gene expression in knockin cells by Western blot. (B) Clonogenic assay results for control and TFAP2B overexpression groups treated with various concentrations of cisplatin. (C) Analysis of clonogenic assay results. n.s. indicates *P* > 0.05, * indicates *P* ≤ 0.05, *** indicates *P* ≤ 0.001. (D) IC50 measurement of cisplatin in cells overexpressing TFAP2B.

## 4. Discussion

This study addresses the significant challenge of platinum resistance in ovarian cancer by identifying genetic markers, constructing a predictive resistance model, and elucidating underlying molecular mechanisms. From the TCGA-OV dataset, 123 resistance-related genes were identified, with 15 significantly impacting survival. The predictive model, based on these 15 hub genes, demonstrated high accuracy (AUC = 0.901), surpassing existing models. Key findings include a lower tumor mutational burden and distinct methylation patterns in resistant groups, activation of specific pathways like MYC targets V2 and Oxidative Phosphorylation, and unique intercellular communication patterns in single-cell analysis. Among the top genes, TFAP2B was highlighted for its significant role in modulating cisplatin resistance.

The discovery of 15 genes associated with platinum resistance and overall survival aligns with and expands upon recent findings in the field. For instance, studies by Cohen YC et al. identified key gene FCRL5 implicated in cancer resistance mechanisms, emphasizing the role of tumor heterogeneity, immune microenvironment [25]. And Yang Q et al. also reported TFAP2B, a UCHL3-related prognostic signature, may be associated with platinum resistance in ovarian cancer [26]. More importantly, in our study among the 15 core genes associated with drug resistance and prognosis, TFAP2B emerged as a significant contributor, underscoring its potential as a therapeutic target. The overexpression of TFAP2B in A2780 cells increased sensitivity to cisplatin, while in A2780cis cells, it inhibited cell growth, indicating TFAP2B’s dual role in modulating resistance. This finding is supported by studies demonstrating the involvement of transcription factors like TFAP2B in regulating drug resistance mechanisms [26].

The PRSM’s predictive capability is particularly noteworthy, offering an improvement over previous models by Zheng et al. (2020) and Li et al. (2021), who reported AUCs ranging from 0.75 to 0.85 in similar contexts [22, 23]. Besides, based on this PRSM model, this study unveils the intricate dynamics of chemotherapy resistance, emphasizing the pivotal role of genomic alterations. Specifically, our analysis of Copy Number Variation (CNV) reveals a notable decrease in the percentage of CNV mutations within the cisplatin-resistant cohort when juxtaposed with their cisplatin-sensitive counterparts, correlating with a broader pattern of genomic stability in the resistant group [27]. Additionally, a significant reduction in Tumor Mutational Burden (TMB) is observed in the resistant group, suggesting a lower probability of forming immunogenic neoantigens that could potentially enhance the efficacy of immune-based therapies [28]. Concurrently, the Homologous Recombination Deficiency (HRD) scores, serving as a proxy for a tumor’s capability to repair DNA double-strand breaks through homologous recombination, are substantially lower in the resistant group. This suggests an altered DNA repair mechanism potentially contributing to cisplatin resistance [29]. Moreover, the load of Single Nucleotide Variant (SNV) Neoantigens is significantly reduced in the resistant group, indicating fewer novel antigens for immune recognition and response, with implications for the effectiveness of immunotherapies in overcoming drug resistance [30]. Together, these findings underscore a complex interplay of genomic stability and immune evasion mechanisms that underpin cisplatin resistance, offering insights into potential therapeutic targets and prognostic markers [31].

Pathway feature analysis revealed significant enrichment of specific pathways in the platinum-resistant group, such as Myc targets and oxidative phosphorylation signaling, aligning with research by Reyes-González JM et al. [32]and Zhao Z et al. [33]that implicated these pathways in cancer chemoresistance. The MYC targets V2 pathway, integral to cell cycle regulation, proliferation, and apoptosis, exhibits enhanced activity in resistant cells, suggesting a pivotal role in promoting an aggressive cancer phenotype that can effectively evade cisplatin’s cytotoxic effects [34]. This upregulation indicates a shift towards increased cellular proliferation and survival, enabling cancer cells to withstand the DNA damage inflicted by cisplatin treatment [35]. Simultaneously, the augmented activity of the Oxidative Phosphorylation pathway in resistant cells underscores a metabolic adaptation conducive to sustaining energy production under the stress of chemotherapy[36]. This elevation in metabolic efficiency is critical for cancer cells to maintain their viability and continue proliferating despite the presence of cisplatin. The enhancement of Oxidative Phosphorylation not only supports the increased energetic demands of resistant cancer cells but also may contribute to the maintenance of redox balance, further facilitating resistance to cisplatin-induced oxidative stress [33]. These pathway alterations in resistance group may reveal a complex adaptive response by cancer cells to overcome cisplatin therapy, highlighting potential therapeutic targets to counteract resistance and improve treatment outcomes.

In our single-cell analysis, the presence of both resistant and sensitive cancer cells within the same ovarian cancer (OV) sample provides insight into the phenomenon whereby patients initially responsive to platinum therapy may subsequently develop secondary resistance. This observation aligns with the findings of Zou et al. [37], who noted that complete responses to drug therapies in tumors are uncommon, with therapy affecting only certain subpopulations within a tumor. Our analysis further demonstrated that the majority of the 15 hub genes were expressed not only in epithelial (cancer) cells but also in NK cells, suggesting that these genes may play a pivotal role in the tumor stroma and epithelial interactions, potentially dictating the response to platinum therapy. Moreover, the analysis highlighted the involvement of specific oncogenic genes and pathways, such as the PRSS1 gene and the WNT signaling pathway, in cell communication within the resistant group. The WNT signaling pathway has been previously identified as a significant contributor to cisplatin resistance in OV [38, 39], while the activation of PRSS1 has been implicated in promoting resistance to cisplatin among ovarian cancer patients [40]. These findings underscore the complex interplay of cellular and molecular mechanisms driving platinum resistance in ovarian cancer, emphasizing the need for targeted approaches to overcome this challenge.

The study’s most significant academic contributions include the development of a robust predictive model for platinum resistance, which offers superior accuracy compared to existing models. This model not only enhances our understanding of genetic and epigenetic factors contributing to resistance but also provides a practical tool for predicting treatment responses. The identification of TFAP2B as a key modulator of cisplatin resistance represents a potential breakthrough in targeting drug-resistant ovarian cancer. Additionally, the integration of single-cell analysis offers a comprehensive view of the cellular heterogeneity and intercellular interactions involved in resistance, contributing to the broader understanding of the tumor microenvironment’s role in drug resistance.

Despite these advancements, the study has limitations. The predictive model, while accurate, requires validation in larger, more diverse cohorts to ensure its generalizability. The study primarily focuses on genetic and epigenetic factors, suggesting a need for further research into other potential mechanisms of resistance, such as proteomic and metabolomic changes. Additionally, while TFAP2B’s role is significant, its exact mechanisms of action in resistance pathways need to be elucidated through in-depth functional studies. Future research should also explore the therapeutic potential of targeting TFAP2B and other identified genes in clinical settings to develop effective treatments for platinum-resistant ovarian cancer.

## 5. Conclusions

We conducted a comprehensive analysis of the TCGA-OV dataset to develop a platinum resistance scoring model (PRSM) for predicting patient survival and platinum chemotherapy response in ovarian cancer. We explored mutations, pathways, and epigenetic landscapes, revealing novel targets and regulatory mechanisms for overcoming platinum resistance. Additionally, we identified a core regulatory gene, TFAP2B, which plays a pivotal role in cisplatin resistance.

## 6. List of abbreviations

PRSM: the platinum resistance scoring model
OV: ovarian cancer
TCGA: The Cancer Genome Atlas
GEO: Gene Expression Omnibus
MSigDB: Molecular Signatures Database
DEGs: Differentially Expressed Genes
ROC: receiver-operating characteristic
OS: overall survival
PFS: progression-free survival
ssGSEA: Single-sample gene set enrichment analysis
FC: fold change
GOBP: Gene Ontology Biological Process
KEGG: Kyoto Encyclopedia of Genes and Genome
CNV: Copy Number Variation
TMB: Tumor Mutational Burden
SNV: Single Nucleotide Variant
HRD: Homologous Recombination Deficiency

## Data accessibility

All data generated or analysed during this study are available from public databases. The detailed information of these data sets is stored in Table 1. R scripts are stored in Supplementary Data Sheet 1.zip.

## Authors’ contributions

The concept of the study was planned by Y.H., W.Q., G.C.; The data processing and analysis were performed by Y.H., and P.W.. Experiments were conducted, analyzed, and interpreted by J.X., B.Z., L.H., X.L., and H.N.. Y.H. drafted the manuscript. W.Q., and G.C. edited the manuscript and provided valuable suggestions. All authors contributed to the article and approved the submitted version.

## Acknowledgements

The authors would like to thank TCGA, GEO and Mendeley data projects for data sharing.

## Supporting information

Additional file 3:

Supplementary Table S1. List of Signature Genes in TCGA Cox pro-portional hazard regression models.

Additional file 4:

Supplementary Table S2. List of lncRNA-miRNA-mRNA pairs.

Additional file 5:

Supplementary Table S3. network correlations analysis.

Additional file 6:

Supplementary file 1 : uncropped Gels and Blots image(s)

Additional file 7:

Supplementary Data Sheet 1.zip: R script.

---

**Fig. S2.**
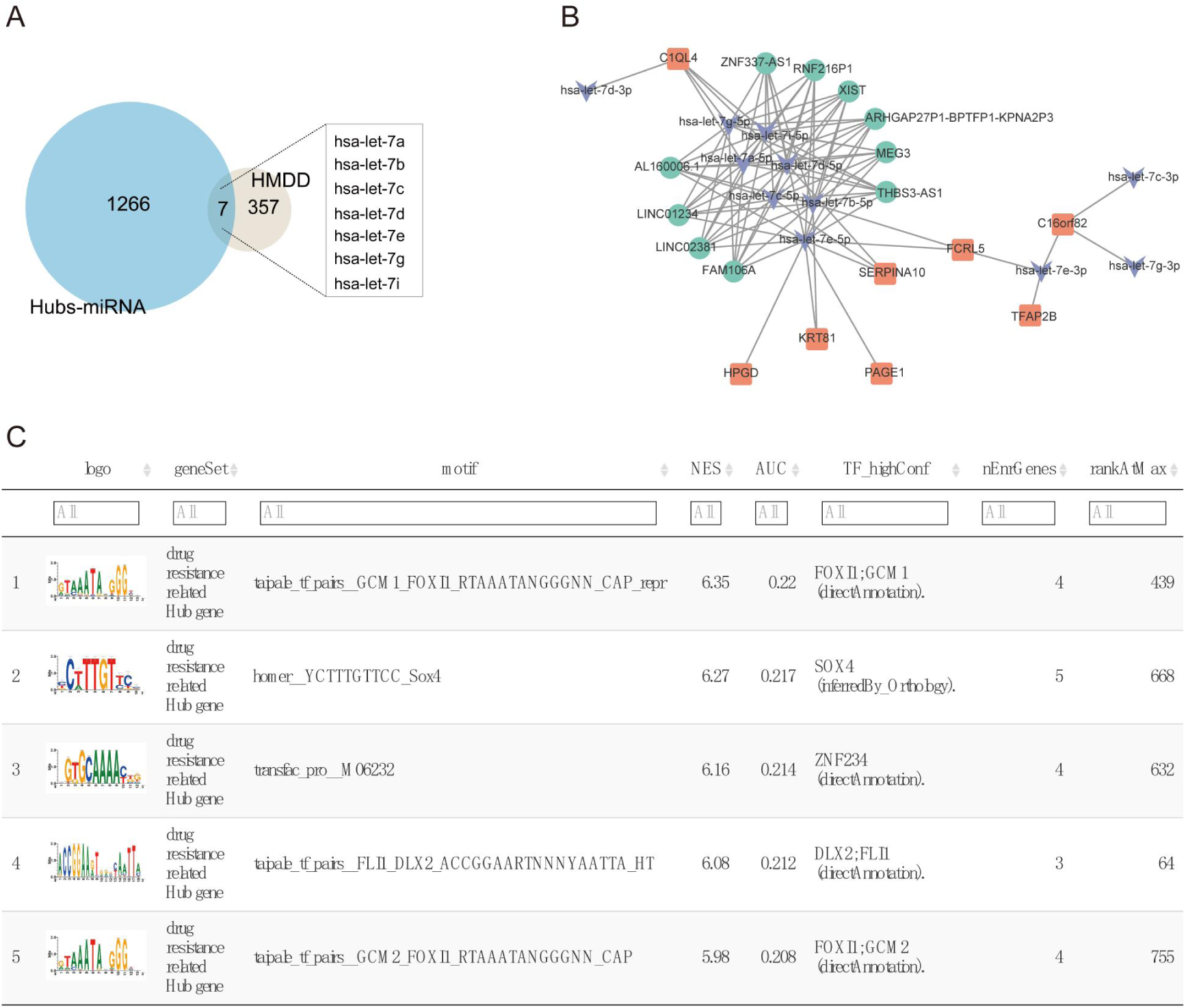
Analysis of the ceRNA Network and TF Regulatory Network Related to Chemotherapy Resistance Hub Genes. A. Venn diagram of ovarian cancer-associated miRNAs and miRNAs related to 15 hub genes extracted from the miRWalk database. B. The ceRNA network of hub genes, with orange-red representing hub genes, purple for miRNAs, and green dots for LncRNAs. C. Enrichment analysis of transcription factors for hub genes.

