## Supplementary figures and images for "A 15-Gene prognostic signature with TFAP2B functioning in Platinum Resistance of Ovarian Carcinoma"

### Supplementary file1

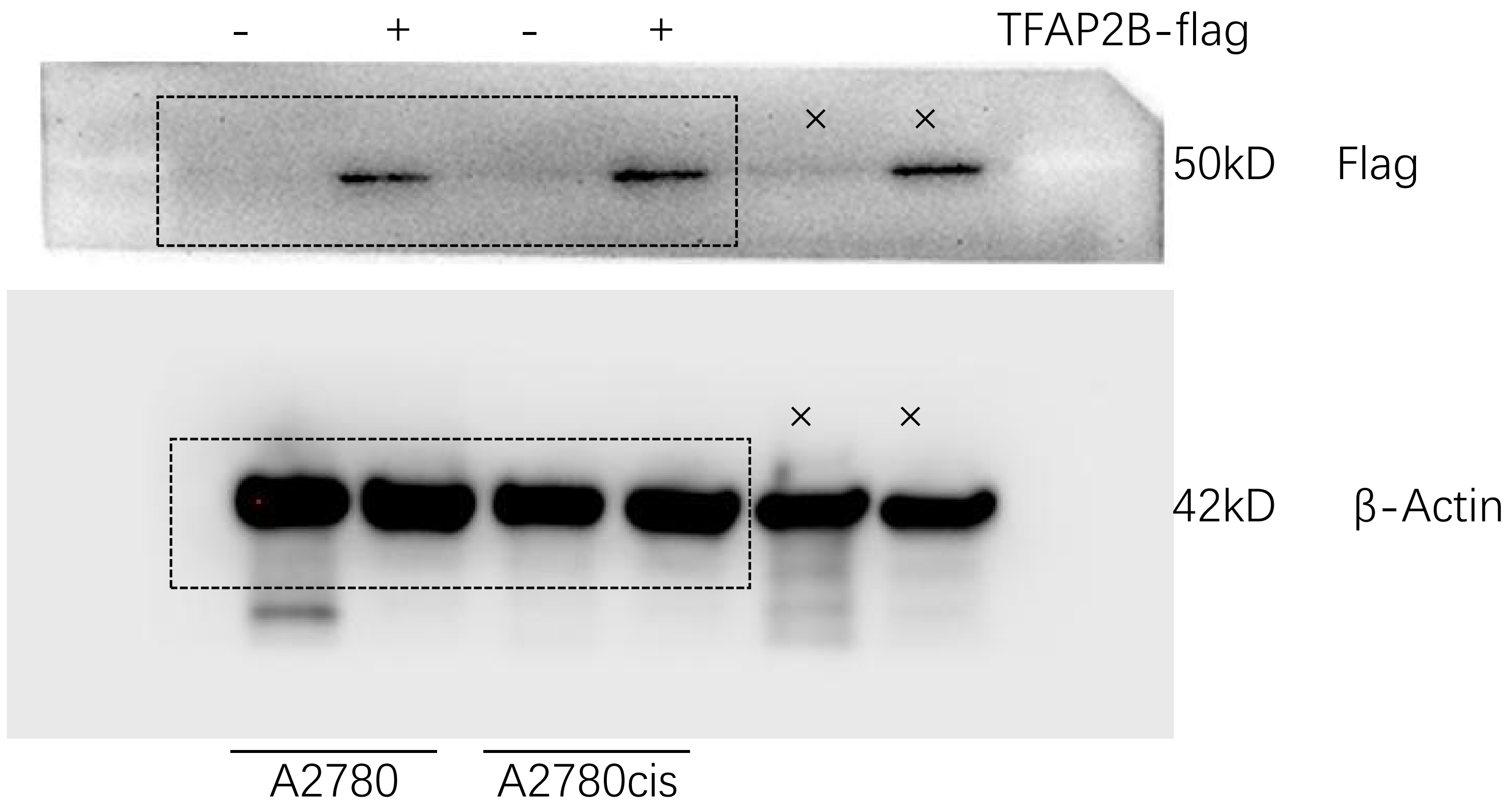
